# Combination of secondary plant metabolites and micronutrients against Alzheimer disease in a SH-SY5Y-APP695 cell model

**DOI:** 10.1101/2023.04.24.538048

**Authors:** Lukas Babylon, Julia Meißner, Gunter P. Eckert

## Abstract

Alzheimer’s disease (AD) is characterized by mitochondrial dysfunction, increased Aβ levels and altered glycolysis. So far, there is no cure for AD, therefore it is important to take preventive or supportive action against AD. The cocktail (SC) tested in this study consists of the substances hesperetin (HstP), magnesium-orotate (MgOr) and folic acid (Fol), as well as the combination (KCC) of caffeine (Cof), kahweol (KW) and cafestol (CF). All the compounds showed positive results in the above mentioned fields of AD. The question arose whether a combination of all of them would also positively affect all three fields of AD. In this regard, SH-SY5Y-APP_695_ cells were incubated with SC and ATP levels, complex respiration, Aβ levels, ROS levels, lactate and pyruvate levels were examined. The SC increased the endogenous respiration of the cells while significantly decreasing the Aβ^1-40^ levels. SC has no significant effects on the other parameters. In summary, the combination of all compounds did not show the desired success that we hoped for, but the cocktail has potential to be further investigated. It is possible that the results will improve by changing the combinations or by adjusting the concentrations.

## 1. Introduction

Alzheimer disease (AD) has now been placed on the WHO’s list of global health priorities. Since its first description in 1906, there is still no effective treatment against it [1]. In 2018, 1.73% of the population in the European Union was living with dementia and this figure is expected to reach 2.0% by 2050 [2]. AD is the most common form of dementia and is on the rise due to an aging population [3]. AD is characterized by a progressive loss of memory, speech, personality and behavior, which is accompanied by limitations in the quality of life [3]. Signs including the presence of amyloid beta (Aβ) which accumulates extracellularly leading to plaque formation and hyperphosphorylated tau characterize AD [4, 5]. Furthermore, AD leads to impaired glucose metabolism [6] and increased reactive oxygen species (ROS) production, which in turn leads to increased oxidative stress and its consequences [7, 8]. In addition, another event that occurs at the onset of the disease is mitochondrial dysfunction (MD) [9]. This leads to reduced metabolism, increased ROS, lipid peroxidation and finally to apoptosis [9, 10]. The first sign of MD is a decrease of enzymes from the oxidative metabolism [11, 12] leading to a limited function of the electron transport chain (ETC) and thus to a reduced complex activity of complexes I and IV. This in turn leads to a lower membrane potential and to lower ATP levels [13, 14]. These defects lead to increased ROS production and defects within the mitochondrion, which contributes to an increase in mitochondrial dysfunction [15]. Since there is no therapy for AD, it is important to intervene early in the disease process to delay possible symptoms as long as possible and to treat and compensate for the defects as well as possible.

One way to support the various defects in AD is through the use of secondary plant compounds, especially flavonoids [16–18]. One such flavonoid is hesperetin (HstP). HstP is an aglycon of hesperedin and is mainly found in the peel of oranges and other citrus fruit [19]. Its possible ability to cross the blood-brain barrier makes it a potential candidate for neurodegenerative diseases [20]. HstP was shown to have a protective effect on neurons previously damaged by Aβ [21]. Hesperetin improve cognitive abilities by increasing BDNF levels in rats. Furthermore, it increases the activities of CAT, SOD, GRX and GPX and cytoprotective effects of hesperetin against Aβ-induced impairment of glucose transport have been reported [22]. Another possibility would be treatment with biofactors [23], defined as substances that the body needs for normal physiological functioning and/or that have health-promoting and/or disease-preventing biological activities, and that can interfere with pathophysiological processes leading to AD [24–26]. Such biofoctors are magnesium-orotate (MgOr) and folic acid (Fol). Folate is an important product in metabolism, when deficient it results in insufficient DNA and also mtDNA synthesis and stability, which is why it can lead to oxidative stress, a process associated with AD. This leads to neuronal aggravation and increased cell death. A deficiency leads to impaired methylation of enzymes and promoter regions of genes involved in AD [27, 28]. MgOr is the magnesium salt of the Ortic acid and a good source of Mg for deficiencies. Ortic acid is a key substance in biosynthetic pathway as well as in the synthesis of glycogen and ATP [29]. Mg is involved in many synthesis processes, as well as in maintaining physiological nerve and muscle function [30–32]. Many different mechanisms related to Mg and AD are discussed. It is thought to positively influence various inflammatory pathways and to influence APP processing towards alpah-secretase, as well as preventing Aß from crossing the blood-brain barrier on a large scale [33, 31]. Another product discussed in connection with AD is coffee, or rather its ingredients. Here in particular caffeine, kahweol and cafestol. Coffee is one of the most commonly consumed beverages [34]. Coffee is supposed to have cardio-, hepa- and neuroprotective effects [35]. The consumption of caffeine and coffee is associated with a lower risk of AD [36–38]. Data on kahweol and cafestol are scarce, but there is evidence that both have neuroprotective effects [39–41].

The substances listed here have been tested by us in various experiments in an early cell model of AD. All substances improve the symptoms triggered by AD in different areas of AD. HstP improves mitochondrial dysfunction [42], MgOr and Fol reduce Aβ levels within cells [43] and caffeine, kahwol and cafestol (KCC) affects glycolysis [44]. In this study, we asked ourselves whether a combination of all substances leads to an improvement in all three areas of AD. For this purpose, a cocktail (SC) of all compounds was used and tested in a SH-SY5Y-APP_695_ cell model.

## 2. Material and Methods

### 2.1 Cell culture

Human SH-SY5Y-APP_695_ cells were incubated at 37°C in DMEM in an atmosphere 5% CO_2_. DMEM was mixed with 10% heat inactivated fetal calf serum, streptomycin, penicillin, hygromycin, non-essential amino acids and sodium pyruvate. The cells were split on average every 3 days and used for experiments when they reached 70-80% growth.

### 2.2 Cell treatment

Cells were incubated with a cocktail for 24 h each unless indicated. This cocktail contains a mixture of hesperetin 10μM (HstP), magnesium orotate 200μM (MgOr), folic acid 10μM (Fol), caffeine 50μM (Cof), cafestol 1μM (CF) and kahweol 1μM (KW) (together SC). The control (CTR) was a mixture of solvents in which the substances were dissolved ethanol, DMSO and NaOH.

### 2.3 ATP Assay

To determine ATP levels, an ATP luciferase assay was performed. Light is generated by ATP and luciferin in the presence of luciferase. The test was performed using the ATPlite Luminescence Assay System (PerkinElmer, Rodgau, Germany).

### 2.4 Cellular respiration

Respiration in SH-SY5Y_695_ cells was measured using an Oxygraph-2k (Oroboros, Innsbruck, Austria) and DatLab 7.0.0.2. Cells were treated according to a complex protocol developed by Dr. Erich Gnaiger [45]. They were incubated with different substrates, inhibitors and uncouplers. First, cells were washed with PBS (containing potassium chloride 26.6 mM, potassium phosphate monobasic 14.705 mM, sodium chloride 1379.31 mM, and sodium phosphate dibasic 80.59 mM) and scraped into mitochondrial respiratory medium (MiRO5) developed by Oroboros [45]. Cells were then centrifuged, resuspended in MiRO5 and diluted to 106 cells/ml. After 2 ml of cell suspension was added to each chamber and endogenous respiration was stabilized, cells were treated with digitonin (10 μg/106 cells) to permeabilize the membrane, leaving the outer and inner mitochondrial membranes intact. OXPHOS was measured by addition of complex I and II substrates malate (2 mM), glutamate (10 mM), and ADP (2 mM) followed by succinate (10 mM). The stepwise addition of carbonyl cyanide-4-, before being evaluated by measuring R123 fluorescence (trifluoromethoxy) phenylhydrazone, showed the maximum capacity of the electron transfer system. Rotenone (0.1 mM), a complex I inhibitor, was injected to measure complex II activity. Oligomycin (2 μg/mL) was then added to measure leak respiration. Inhibition of complex III by addition of antimycin A (2.5 μM) was used to determine residual oxygen consumption, which was subtracted from all respiration states. The activity of complex IV was measured by addition of N, N, N′-tetramethyl-p-phenylenediamine (0.5 mM) and ascorbate (2 mM). To measure the sodium autooxidation rate, azide (≥ 100 mM) was added. Subsequently, complex IV respiration was corrected for autoxidation.

### 2.5 Citrate Synthase Activity

Cell samples from respirometry were frozen away at -80°C for citrate synthase determination. Samples were spiked and mixed with the reaction mix (0.31 mM acetyl coenzyme A, 0.1 mM 5,5′-dithiol-bis-(2-nitrobenzoic acid) (DTNB), 50 μM EDTA, 0.5 mM oxaloacetate, 5 mM triethanolamine hydrochloride and 0.1 M Tris-HCl) and warmed to 30°C for 5 minutes. Samples of 40μl each were applied as a triplet and 110μl of reaction mix was added and the absorbance was measured.

### 2.6 Protein quantification

To determine the protein content of the samples, a PierceTM Protein Assay Kit (Thermo Fisher Scientific, Waltham, MA, USA) was used and the experiment was performed according to the manufacturer’s instructions, using bovine serum albumin as the standard.

### 2.7 Aβ_1-40_ measurement

After 24h incubation, Aβ_1-40_ was determined using an HTRF amyloid beta 1-40 kit (Cisbio, Codolet, France). The protocol has been described previously [42]. Aβ_1-40_ levels were normalized against protein levels.

### 2.8 ROS quantification

A DCFDA/H2DCFDA kit was used to determine ROS levels (ab113851; Abcam, Cambridge, UK). Cells were seeded in 96 well plates and incubated for 24h. The experiment was then performed according to the manufacturer’s instructions.

### 2.9 Lactate and pyruvate measurement

Frozen cell samples, which were previously incubated for 24h, were thawed at room temperature. An assay kit (MAK071, Sigma Aldrich, Darmstadt, Germany) was used to determine pyruvate levels. For the determination of lactate values, a lactate assay kit was used (MAK064, Sigma Aldrich, Darmstadt, Germany). The values were normalized to the protein content.

### 2.10 Statistics

Unless otherwise stated, values are presented as mean ± standard error of the mean (SEM). Statistical significance was defined for *p* values * p < 0.05, ** p < 0.01, *** p < 0.001 and **** p < 0.0001. Statistical analyses were performed by applying student’s unpaired t-test and one-way ANOVA with Tukey’s multiple comparison post-hock test (Prism 9.5 GraphPad Software, San Diego, CA, USA).

## 3. Results

### 3.1 Overview

The SC used in this study is a combination of different single substances, which have been previously tested as single substances or combinations in previous studies. This table lists the individual substances and the results of the SC.

**Table 1:** Overview of the experiments discussed in this paper and their respective effects.

|  | HstP | MgOr + Fol | Cof + KW + CF | SC |
| --- | --- | --- | --- | --- |
| ATP level | ↑ | = | ↑ | ↑ |
| OXPHOS | = | = | = | ↑ |
| ROS | ↓ | = | = | = |
| Aβ level | = | ↓ | = | ↓ |
| Lactate | = | ↓ | = | = |
| Pyruvate | = | = | ↑ | = |
| L/P Ratio | = | = | ↓ | = |
| Source | [42] | [43] | [44] |  |

↑ = significant increase; ↓ = significant decrease; = no significant change; L/P = lactate-pyruvate-ratio

### 3.2 Mitochondrial parameter

First, we examined the cells for various mitochondrial functions, including ATP levels, respirometry and related citrate synthase activity. Incubation with SC resulted in a significant increase in ATP levels compared to control (p = 0.0031, Fig. 1A). Since the increased ATP levels may be due to a change in respiratory chain activity, this was investigated next. A significant increase in endogenous respiration of SC and SCTR was found (p = 0.0314). In addition, there were no significant changes in SC versus SCTR. Except for complex II and leak II, the O2 consumption of SC tended to be higher than that of SCTR (Fig. 1B). There was no difference in citrate synthase activity between the groups (Fig. 1C).

**Figure 1:**
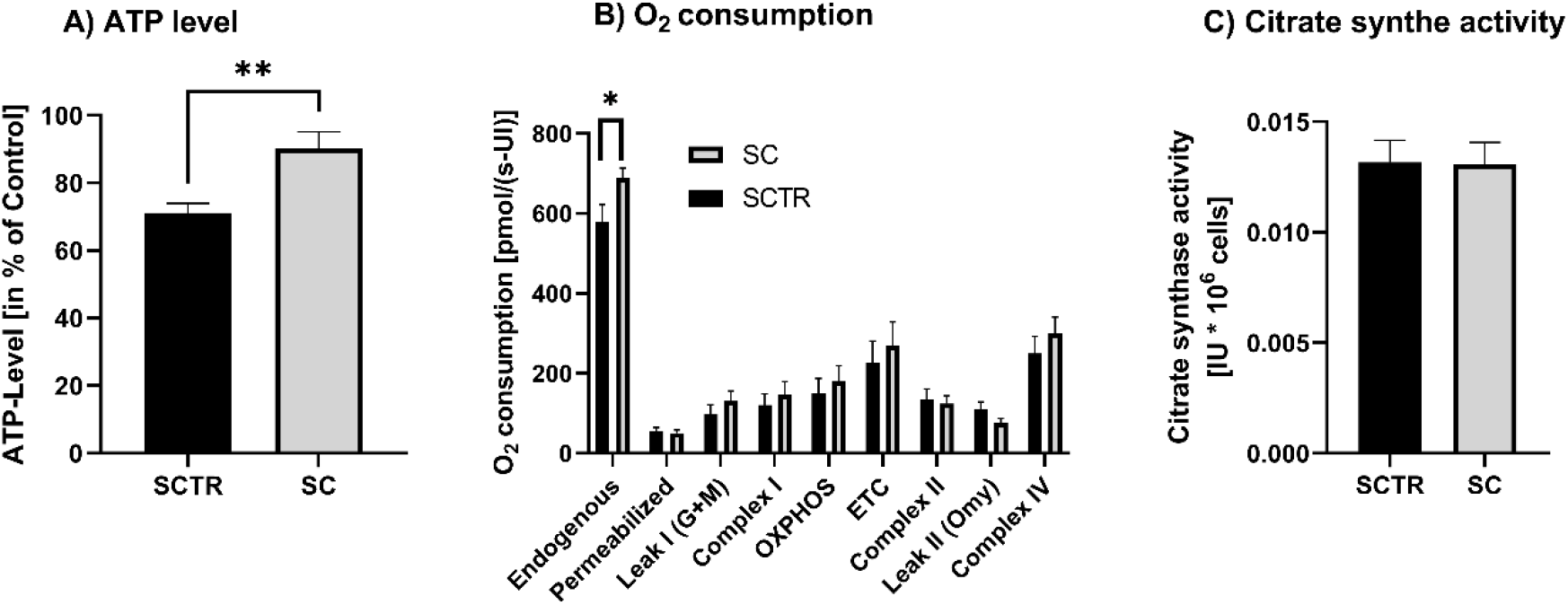
ATP level, respiration and citrate synthase activity of SH-SY5Y-APP_695_ cells incubated for 24h with SC. (A) ATP level of SH-SY5Y-APP_695_ cells incubated with SC. Cells treated with cell culture medium served as control (100%). N = 11. (B) Respiration of SH-SY5Y-APP_695_ cells incubated with SC compared to the control. SH-SY5Y-APP_695_ cells adjusted to international units (IU) of citrate synthase activity. N = 16. (C) Citrate synthase activity of SH-SY5Y-APP_695_ cells incubated with SC compared to control. N = 16. Significance was determined by Student’s unpaired t-test. Data are displayed as the mean ± SEM. *P > 0.05, ***p> 0.001. SC = cocktail, SCTR = control.

### 3.3 Aβ_1-40_ level

To investigate the Aβlevels, the cells were incubated with SC or SCTR for 24 h (Fig. 2). SC significantly decreased the Aβlevels of the cells compared to the control (p = 0.0217).

**Figure 2:**
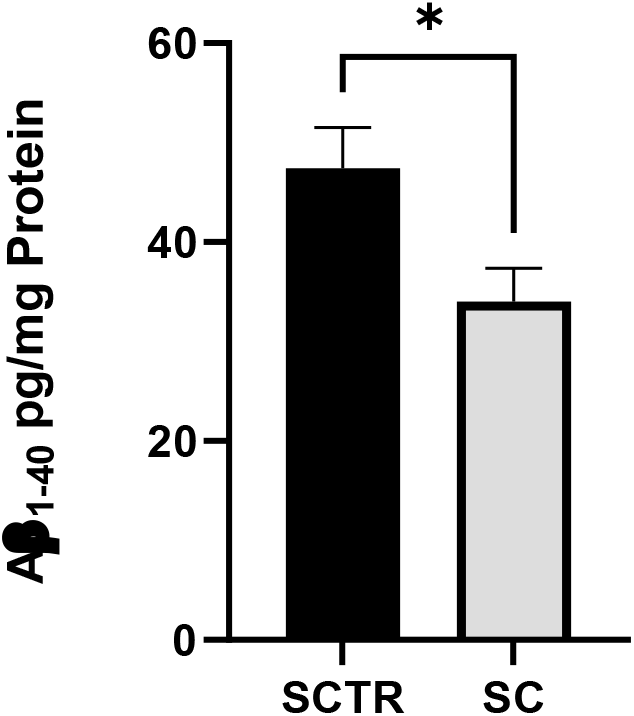
Effect of SC on the Aβ_1-40_ level of SH-SY5Y-APP_695_ cells incubated for 24h. N = 9. Aβ_1-40_ levels were adjusted to the protein content. Significance was determined by Student’s unpaired t-test. Data are displayed as the mean ± SEM. *P > 0.05 SC = cocktail, SCTR = control

### 3.4 ROS

To investigate the effects of incubation on ROS levels, cells were incubated with the respective substance for 24 hours (Fig. 3). The SC is based on our previously tested substances, but the measurement of ROS levels by incubation with MgOr, Fol or the combination of both ID63 was missing. This was done here. No significant differences were found.

**Figure 3:**
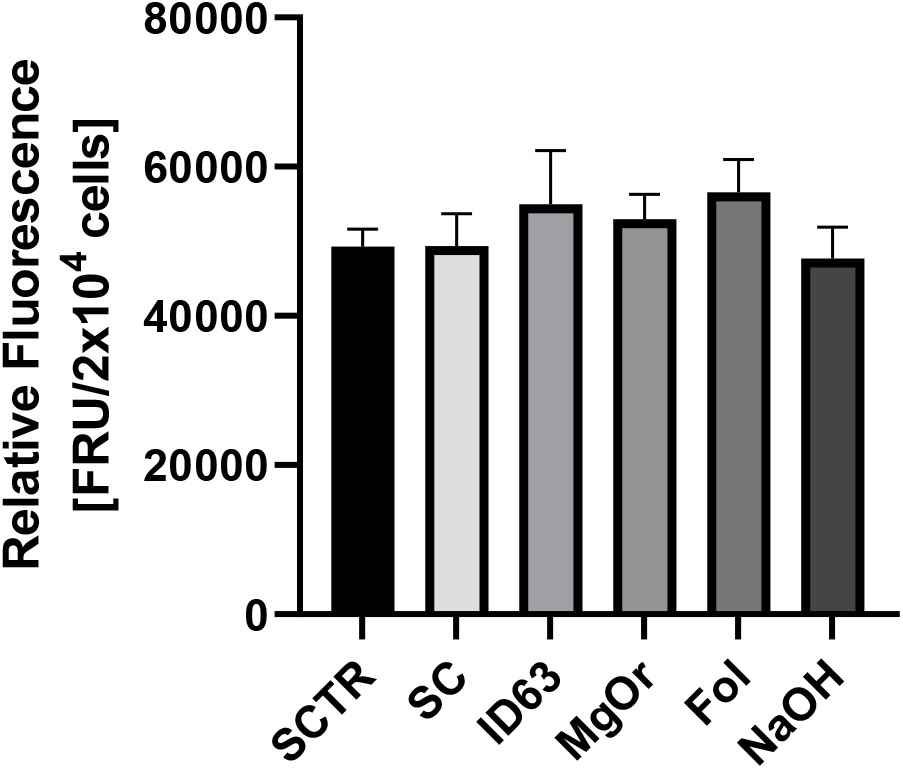
ROS measurement of SH-SY5Y-APP_695_ cells after incubation with KCC for 24h. N = 8. Significance was determined by Student’s unpaired t-test. Data are displayed as the mean ± SEM.

### 3.5 Glycolysis

To investigate the effects of the incubations on glycolysis metabolism, the lactate and pyruvate values and their ratio were determined. First, these values were examined for the cocktail (Fig. 4). There were no significant differences, although the lactate values of the SC tended to be higher than those of the control (p = 0.0983). In contrast, the pyruvate values of the SC tended to be lower than those of the control. This results in a lower ratio for the control (p = 0.0509).

**Figure 4:**
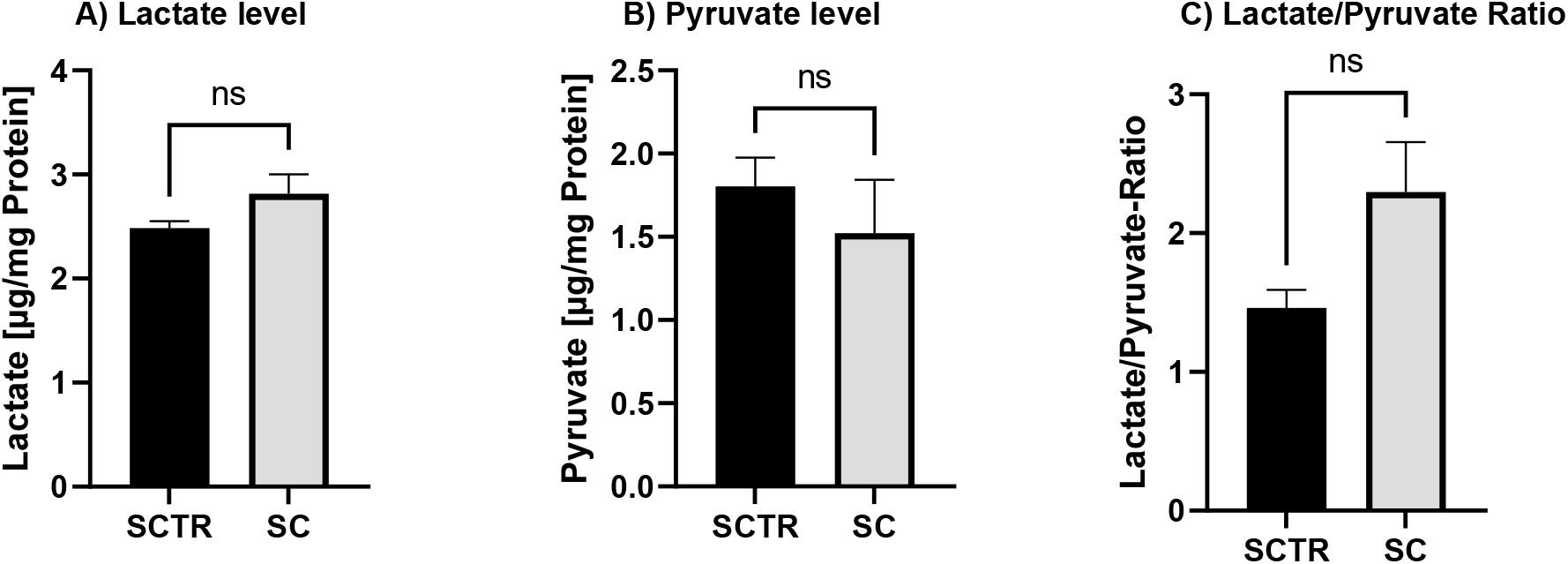
Effect of the on lactate and pyruvate level after 24h of incubation with SC or the control. (A) Lactate level of SH-SY5Y-APP_695_ cells after the incubation compared to the control. (B) Pyruvate level of SH-SY5Y-APP_695_ cells after the incubation with KCC compared to the control. (C) Lactate to pyruvate ratio. N = 10. Levels were adjusted to the protein content. Significance was determined by Student’s unpaired t-test. ns = not significant. Data are displayed as ±SEM.

The lacatet, pyruvate and the resulting ratio to each other were also determined for HstP (Fig 5). We also lacked these data for glycolysis, which is why they were made up for in this framework. Here, no significant differences were found for lactate but pyruvate level were slightly elevated and the ratio were slightly reduced in favor of HstP.

**Figure 5:**
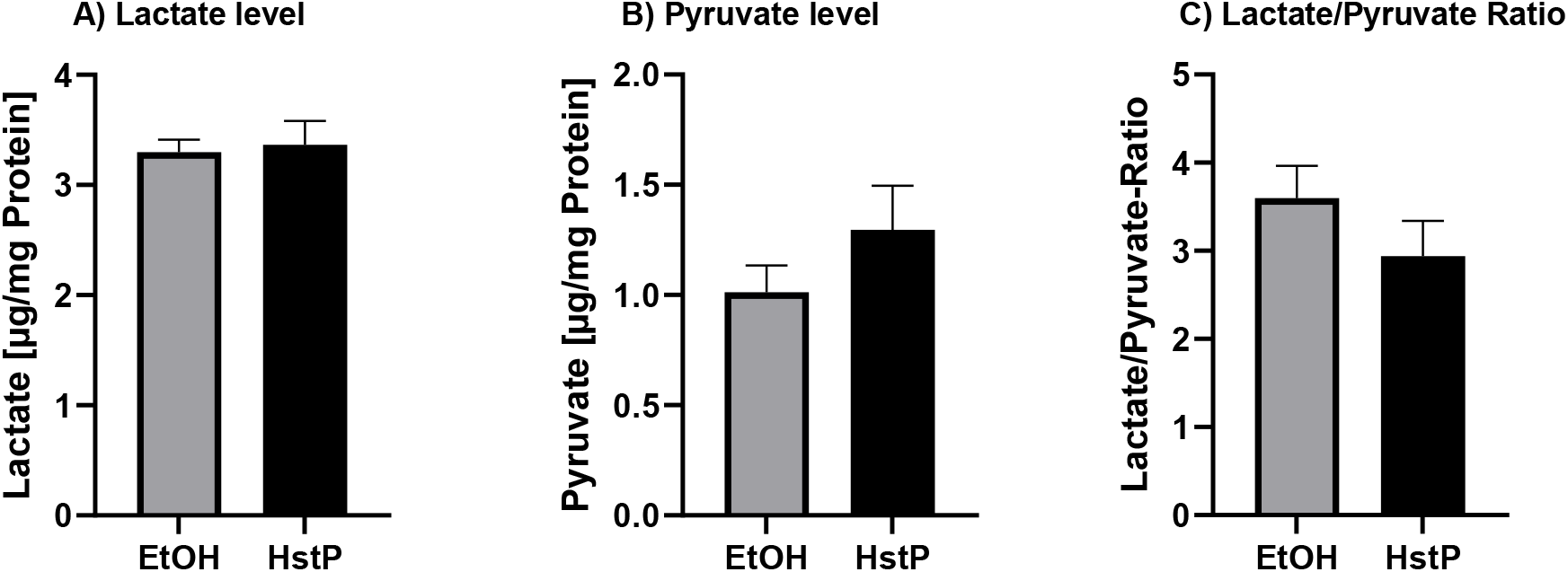
Effect of the on lactate and pyruvate level after 24h of incubation with HstP or the control. (A) Lactate level of SH-SY5Y-APP_695_ cells after the incubation compared to the control. (B) Pyruvate level of SH-SY5Y-APP_695_ cells after the incubation with KCC compared to the control. (C) Lactate to pyruvate ratio. N = 8. Levels were adjusted to the protein content. Significance was determined by Student’s unpaired t-test. Data are displayed as ±SEM.

## 4. Discussion

Mitochondrial dysfunction is one of the core issues in the development of AD, as described earlier and is characterized by a decrease in ATP [13, 14, 46]. The SC increased the ATP levels when compared to the control (Fig. 1A). However, the values did not reach those of the medium control, which suggests that the large number of substances and solvents used behaved in a contrainduced manner, which was, compensated for in comparison with the SC control. This is possibly due to the individual substances. To understand the effects of the SC, we need to look at the influence of the individual substances on the various parameters in the cell. HstP caused ATP levels to increase in our studies as well as in *Biesemann et. al* [47]. In the same way, Mg and Fol are involved in the provision of ATP. Deficiencies of these nutrients leads to a decrease in ATP production [27, 31]. These deficits are also evident in AD, in which Mg and Fol levels are found reduced in brain tissues of patients [48–51]. However, our cell model is not one in which there is a deficit of Mg and Fol. Furthermore, in our studies, no increase in ATP levels was observed by the single administration of MgOr and Fol [43]. In contrast, the combination KCC increased ATP levels [source]. Cof alone probably has no effect on ATP levels as shown in our experiments. It is possible that the increase is due to KW and CF [source]. Whereas in the literature the increase of Cof is described [52, 53].

To explain the enhanced ATP levels in some cases, we looked at the OXPHOS after incubation with the SC (Fig. 1B). The OXPHOS is a pathway where ATP is produced, so an increase in the activity of the complexes could result in an increase in ATP [54, 55]. The administration of SC significantly increased endogenous respiration. Furthermore, the respiration of the individual complexes was increased compared to the control reduced. Possible reasons for the increased respiration could be the effects arising from the individual compounds. Activation of the Nrf-2 pathway by HstP would be a conceivable option [56, 57]. Reduction of Nrf-2 results in decreased MMP, decreased ATP levels and impaired mitochondrial respiration [58]. A reduction of Aβ levels by MgOr and Fol, would be also a possible explanation [43]. This could reduce the impairment of complexes by Aβ. In our single study, KCC had no effect on OXPHOS, so the described effect could be due to the other substances [Quelle]. The increased respiration would be a possible reason for the increased ATP levels observed by the combination with the cocktail.

It was interesting for us to see how a combination of all the substances we tested affected ROS levels in SH-SY5Y cells (Fig. 3) and whether the ROS-reducing effect of HstP played a role. [42]. It was shown that the ROS levels of the cocktail did not decreased. Thus, HstP could not play out its potential. It is possible that the amount of substances leads to the fact that no ROS reducing effect has occurred. Fol and MgOr had no effect on ROS levels (Fig. 3), whereas the literature suggests a ROS reducing effect from Mg [59], Fol [60], Cof [61], KW [62] and CF [63]. However, the described effects are difficult to compare because they are either deficit models or models other than our SH-SY5Y cell model were used.

Next, we took a closer look at Aβ levels (Fig. 2). We found that SC significantly lowered Aβ_1-40_ compared to the control. Considering the SC results, there is an additive effect on Aβ_1-40_ levels. Of the compounds we tested, only MgOr and Fol had the potential to significantly lower Aβ levels [43], but in combination as SC, all substances lower Aβ levels. Indeed, all our tested substances show the same potential in the literature [56, 64–66]. The question now is whether the effect we observed is due to the combination of MgOr and Fol alone, or whether all substances are exerting their potential. It is possible that the additive effect results in a suppression of BACE1 and APP expression and a greater reduction in β-secretase activity than we observed with MgOr and Fol. This should be confirmed in further experiments.

As a final point, the cocktail was examined for the effects of glycolysis (Fig. 4). Here, the incubation showed an increase in lactate levels, with a concomitant decrease in pyruvate levels and an L/P in favor of lactate. Based on these data, it can be assumed that the cells tend to an increased anaerobic respiration due to the SC. It is interesting to note that, as with MgOr and Fol, there is an Aβ reducing effect, but this effect does not show up in glycolysis as with MgOr and Fol [43]. It has been shown that cells protect themselves from Aβ by increasing anaerobic respiration, with the help of the Warburg effect [67–69]. This manifests itself in an increase in anaerobic respiration and an increase in lactate [70], as in our case. At the same time, OXPHOS is reduced as the cells switch from aerobic to anaerobic respiration. This effect does not occur in our case because OXPHOS is increased. Furthermore, the Aβ levels are decreased by the SC and thus the Warburg effect is no longer necessary for the cells. It is possible that the increase in OXPHOS is not sufficient for the cells to synthesize enough ATP, which is why anaerobic respiration must be increased at the same time to synthesize sufficient ATP. This contributes to the significant increase in ATP levels.

## 5. Conclusion

Each of our substances or combination of substances addressed a different impaired field in AD. HstP affected enhancement MD, MgOr and Fol lowered Aβ levels, and KCC affected glycolysis. Our question was whether a combination of all covered the same three fields. Here, we found that the cocktail only increased ATP levels as well as decreased Aβ levels. Possibly the number of substances had a negative effect on the cells and thus did not achieve the same effects or the substances interfered with each other and hindered their effect. Nevertheless, our studies have shown the potential to combat early AD and therefore require further research. Possibly a reduced number of substances would show a stronger effect. However, this is outside the scope of this study and needs to be investigated elsewhere.

## Author Contributions

investigation, L.B, J.M.; writing—original draft preparation, L.B.; supervision, G.P.E. All authors have read and agreed to the published version of the manuscript.

### Funding

This research received no external funding

### Data Availability Statement

The data presented in this study are available on request from the corresponding author.

